# Frameshifts in Tandem Repeats: Consequences on Protein Physicochemical Properties and Function

**DOI:** 10.1101/2024.06.02.597034

**Authors:** Zarifa Osmanli, Gudrun Aldrian, Jeremy Leclercq, Theo Falgarone, Santiago M. Gomez Bergna, Denis N. Prada Gori, Andrew V. Oleinikov, Ilham Shahmuradov, Andrey V. Kajava

## Abstract

The genetic code uses three-nucleotide units to encode each amino acid in proteins. Insertions or deletions of nucleotides not divisible by three shift the reading frames, resulting in significantly different protein sequences. These events are disruptive but can also create variability important for evolution. Previous studies suggest that genetic code and gene sequences evolve to minimize frameshift effects, maintaining similar physicochemical properties to their reference proteins. Here, we focused on tandem repeat sequences, known as frameshift hotspots. Using cutting-edge bioinformatics tools, we compared reference and frameshifted protein sequences within tandem repeats across 50 prokaryotic and eukaryotic proteomes. Our analysis revealed several intriguing sequence-structure-function correlations. We showed that in contrast to the general tendency, frameshifts within these regions, especially with short repeats, lead to significant changes: increased hydrophobicity and arginine content, new aggregation-prone and transmembrane regions. Overall, frameshifts have stronger effects on tandem repeat regions compared to non-repetitive sequences, and therefore can be a primary cause of altered functions, cellular localization, and the development of various pathologies.

## INTRODUCTION

Genetic information flows from DNA through messenger RNA into protein (1). The genetic code, which translates information from the ‘language’ of nucleic acids to that of proteins, follows a triplet nature, where three nucleotides encode a single amino acid. Due to the triplet nature of the genetic code, an insertion or deletion of a number of nucleotides that is not divisible by three can shift the reading frame, resulting in significantly different protein sequences. Besides the frameshifts induced by the indels, a shifting process can also occur on ribosomes when they translate the same mRNA into different polypeptide chains (2-5). This Programmed Ribosomal Frameshifting serves to increase the protein-coding capacity of genomes (6-7) and to downregulate protein translation in all kingdoms of life (8-9). Splicing of exons in different frames also enriches the diversity of functional genes in eukaryotic organisms (10).

The frameshift event, in which a minor nucleotide-level alteration can result in profound protein-level changes, appears to be exceptionally efficient for the generation of a variability required for the process of evolutionary selection. The frameshift mutations are mostly known for their disruptive nature (11). They typically result in significantly modified protein sequences, often with premature stop codons that, in turn, lead to nonfunctional or potentially harmful products (12). Frameshifts are frequently associated with various health disorder variants, including cancer (13). The growth of high-quality genome sequence data and advancements in proteomics and bioinformatics methodologies enable researchers to conduct large-scale comparisons between reference proteins and their frameshift variants. Several such studies suggested that the genetic code may be optimized during evolution to minimize the effects of errors introduced by frameshifting (14-16). Furthermore, it has been demonstrated that several characteristics of protein sequences, including their hydrophobicity profiles and intrinsic disorder profiles, maintain similarity in the corresponding frameshifted sequences. This result suggests that frameshifting could serve as an evolutionary mechanism for generating new proteins with significantly distinct sequences while maintaining similar physicochemical properties to their parent proteins (17).

In this work, we focused on a comparative proteome-wide analysis of reference proteins and their frameshifted sequences containing tandem repeats (TRs). There are several compelling reasons to gain a more profound understanding of the structural and functional implications of frameshifts that can occur within TRs. First, proteins containing TRs are widespread in genomes, appearing in nearly one-third of human proteins, and in as many as half of proteins from *Dictyostelium Discoideum* or *Plasmodium falciparum* (18-19). Second, the frequency of frameshift events has been shown to increase in TRs having short repetitive units of 1 to 3 residues (20-21). As concerns longer repeats, they originate from tandem duplication (22). The duplication, coupled with frameshift mutations, is proposed to serve as a common mechanism for generating functional innovations in proteins (23). Third, TRs, particularly those with short repeats, often exhibit a pronounced bias in amino acid compositions, resulting in a heightened potential for molecular interactions due to the elevated local concentration of specific physicochemical properties such as hydrophobicity, charge or flexibility (24). The occurrence of frameshift mutations within these TRs has the potential to entirely alter their sequences, resulting in markedly different structural and functional characteristics, with the amplitude of these changes being notably higher than in aperiodic protein sequences. Recognizing the significance of this phenomenon and the absence of systematic studies, our comprehensive proteome-wide analysis aims to investigate the potential changes in protein sequence, structure, and function resulting from frameshifts within TRs.

## MATERIALS AND METHODS

### Selection of datasets

We selected coding sequences (CDSs) of 50 well-studied organisms across prokaryotes and eukaryotes from RefSeq and Ensembl databases (25) (Supplementary Figure 1). Then we obtained a non-redundant set of CDS to avoid duplicated sequences coming from RefSeq and Ensembl databases by clustering the sequences with CD-HIT (26). The reference set includes protein sequences obtained from the translation of the CDSs in the main reading frame. The frameshifted set includes amino acid (AA) sequences obtained from the translation of CDS in two frames (-1 and +1). The minimal length of the frameshifted sequence was chosen at 60 AA. As a result, the reference set had 1 032 420 sequences and the frameshifted one 3 005 214 sequences. For further details, see Supplementary Figure S2.

**Figure 1.**
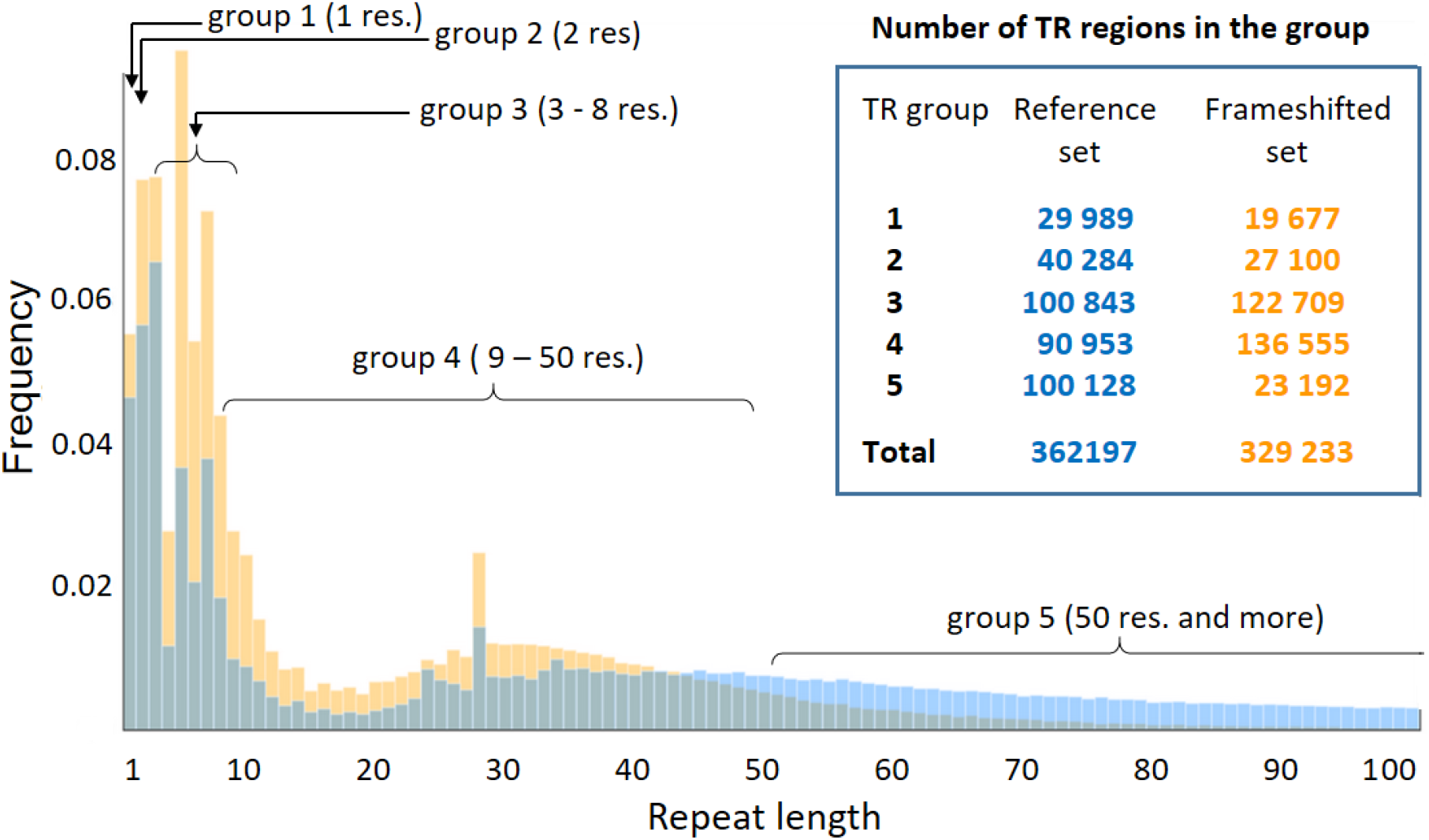
The frequency of TRs depends on their repeat length. The frequencies are computed by dividing the number of TRs with a specific repeat length by the total number of TRs within either the reference (blue) or frameshifted (orange) sets. The histograms depict the frequencies of TRs up to the repeat length of 100 AA. TRs are subdivided into 5 groups based on the length of their repeats, and the inset on the right side of the histogram contains a table showing the number of TRs in each group. In the figure, “res.” stands for AA residues.

### Prediction of structural states of reference and frameshifted sequences

In AA sequences, we predicted regions with Pfam, CATH domains, Short Linear Motifs (SLiMs) by the TAPASS pipeline (27). Aggregation-prone regions (ARs) and exposed aggregation-prone regions (EARs) were predicted by ArchCandy 2.0 (28) and TANGO (29). Intrinsically disordered regions were determined by IUPred (30). The 3D structures were predicted by the AlphaFold2 (AF2) program (31).

### Prediction of TR regions in reference and frameshifted sequences

TRs were predicted by MetaRepeatFinder (MRF), which employed multiple TR finders, each excelling within a specific range of repeat lengths (https://bioinfo.crbm.cnrs.fr/index.php?route=tools&tool=15). In the MRF pipeline, Selectseq and Regex finders predict TRs with repeat units of 1 to 3 AA, T-REKS (32) is the most suitable for the identification of TRs with units till approximately 15 AA in length, and TRUST (33) for repeats longer than 15 residues. A software package pftools, incorporating programs for sensitive generalized sequence profiling (34) was used to detect collagen-like and α-helical coiled coil TRs in protein sequences.

### Clustering of TRs

We used an in-house script for the clustering of TRs. The clustering was based on TR consensus motifs. Consensus sequences of the multiple sequence alignment (MSA) of TRs were retrieved using an algorithm designed for T-REKS (32). The values of each AA frequency were stored in an ordered vector with a constant AA position. These vectors were then utilized as input data for clustering using the DBSCAN algorithm (35).

### Codon usage in different reading frames of homorepeats

We calculated codon usage in the DNA regions where at least 9 AA of the reference reading frame overlap with frameshifted homorepeat regions. For comparison, we also calculated the codon usage in all proteins of the analyzed proteomes. The analysis was done by an in-house script. The following formula was used for computing the frequency of codons that are conserved within TR regions: *Frequency of conserved codons = Number of conserved codons/(number of conserved codons + of variable codons) x 100%*, where codons were considered to be variable if at least one mutation at the DNA level was detected within a TR region.

### AlphaFold2 modeling of conserved structured TR regions from the frameshifted set

We developed in-house scripts to select TR-containing frameshifted sequences for AlphaFold2 (AF2) structural prediction. In the first step, we searched for sequence similarity within frameshifted sets of different studied proteomes from eukaryotes and prokaryotes separately. To find conserved regions of frameshifted sequences, we run the BLAST program with the following parameters: sequence coverage of more than 90%, alignment length >= 30 AA, e-value = 10^−5^, PAM30 as an AA substitution matrix (36). We focused on five proteomes with the highest number of frameshifted sequences (*H*.*sapiens, P*.*troglodytes, C*.*lupus, B*.*taurus, M*.*musculus*). We run the TR-containing frameshifted sequences from each of these proteomes against the remaining 28 frameshifted sets of eukaryotic proteomes. Those frameshifted sequences having regions with at least 1 hit found across 28 eukaryotic species were selected as “conserved” for further analysis. Next, a filter was applied to find the conserved regions, which completely overlap with TR regions. Subsequently, only those sequences that overlapped with structured regions detected by the TAPASS pipeline were selected and then clustered by CD-HIT to eliminate redundancy. Finally, this non-redundant set of the frameshifted sequences was used as input for AF2 modeling by using the Jean-Zay supercomputer at the National Computing Centre for the CNRS (IDRIS, the link to access http://www.idris.fr/eng/info/missions-eng.html) with default parameters. We made a filter to sort out AF2 structural models with ordered regions longer than 20 AA and disordered conformations based on the algorithm used in TAPASS (27). The ordered TR regions of AF2 models having the confidence level plDDT >= 70 were selected for further manual analysis.

## RESULTS

### 1. Distribution of TRs in reference and frameshifted sequences

The periodicity of an AA sequence is encoded in the corresponding DNA sequence. Therefore, if a reference protein contains a TR, its frameshifted sequences also frequently should have TRs in this region. Our analysis supported this conclusion. For example, analysis of *S. sclerotiorum* proteome showed that 33.5% of the TR-containing proteins also have TRs in the other reading frames. Frameshifted AA sequences are usually shorter than the reference proteins due to frequent stop-codons. Therefore, certain TR regions disappear in the frameshifted set.

Analyzing proteomes of 21 prokaryotic and 29 eukaryotic species, we found that 16.60 % of 1 032 420 reference sequences and 9.45 % of 3 005 214 sequences of -1 and +1 frame shifts contain TRs. When counting the TR regions, there are 362 197 regions in the reference set and 329 233 in the -1 and +1 frameshifted sequences (Figure 1). Reference and frameshifted sets have similar dependencies of TR frequencies on repeat lengths with the most frequent TRs concentrated in the region of short repeats (Figure 1). At the same time, the long repeats over 50 residues are rare in the frameshifted sequences in comparison with the reference ones. This can be explained by the shorter total length of the frameshifted sequences compared to the reference ones. The frameshifted TR-containing sequences have an average length of 130 AA with an average repeat length of 19 AA, while the reference sequences average 807 AA with a repeat length of 60 AA. The sequence length should not significantly impact the distribution of the short repeats. Indeed, similarly to the reference set, frameshifted TRs are enriched in the 5- and 7-residue repeats (Figure 1). We also observed a peak at 28-residue repeats in both sets, which in the reference set corresponds to TRs of Zn-finger proteins.

### 2. Subdivision of the analyzed TRs into groups based on the repeat length

Depending on the repeat length, TRs have different structural and functional properties. This interdependence laid the basis for a classification of TRs that form stable 3D structures (37). However, a significant part of TRs are predicted to be unstructured and their probability to be unstructured also depends on the repeat length (38, 19). The shorter the length of the repeat unit, the higher the chance for the TR to be unstructured. Thus, for our analysis it was instrumental to subdivide TRs into several groups depending on the repeat length. We suggest a division that is similar but not identical to the structural classification (37). In the structural classification, the repeat lengths of the Classes are overlapped, while our current analysis required groups with non-overlapped repeat lengths. As a result, we used the following five non-overlapping groups (Figure 1). Group 1 includes homorepeats with the repeat unit of 1 residue. Group 2 consists of TRs with 2-residue repeat units. Both groups belong to Class I of the structural classification. TRs of group 3 with repeat units ranging from 3 to 8 residues belong to Class II and partially Class III in the structured classification. Group 4 has TRs with repeats from 9 to 50 residues. In the structural classification, this group is predominantly composed of Class III TRs, with some inclusion of Class IV and Class V TRs. Group 5 contains TRs with repeats of more than 50 residues. It corresponds to TRs from Class V and partially from Class IV of the structural classification. Figure 1 displays the counts of TRs in each group.

### 3. AA composition of TR regions in the reference and frameshifted sequences

The 3D structures of proteins are determined by their AA sequences. At the same time, the AA composition of proteins also allows us to conclude about their structural states. We compared the AA composition of the TR of the reference and frameshifted sequences. This analysis was done separately for eukaryotic and prokaryotic proteins (Figures 2A and 2C and Supplementary Tables S1 and S2).

**Figure 2.**
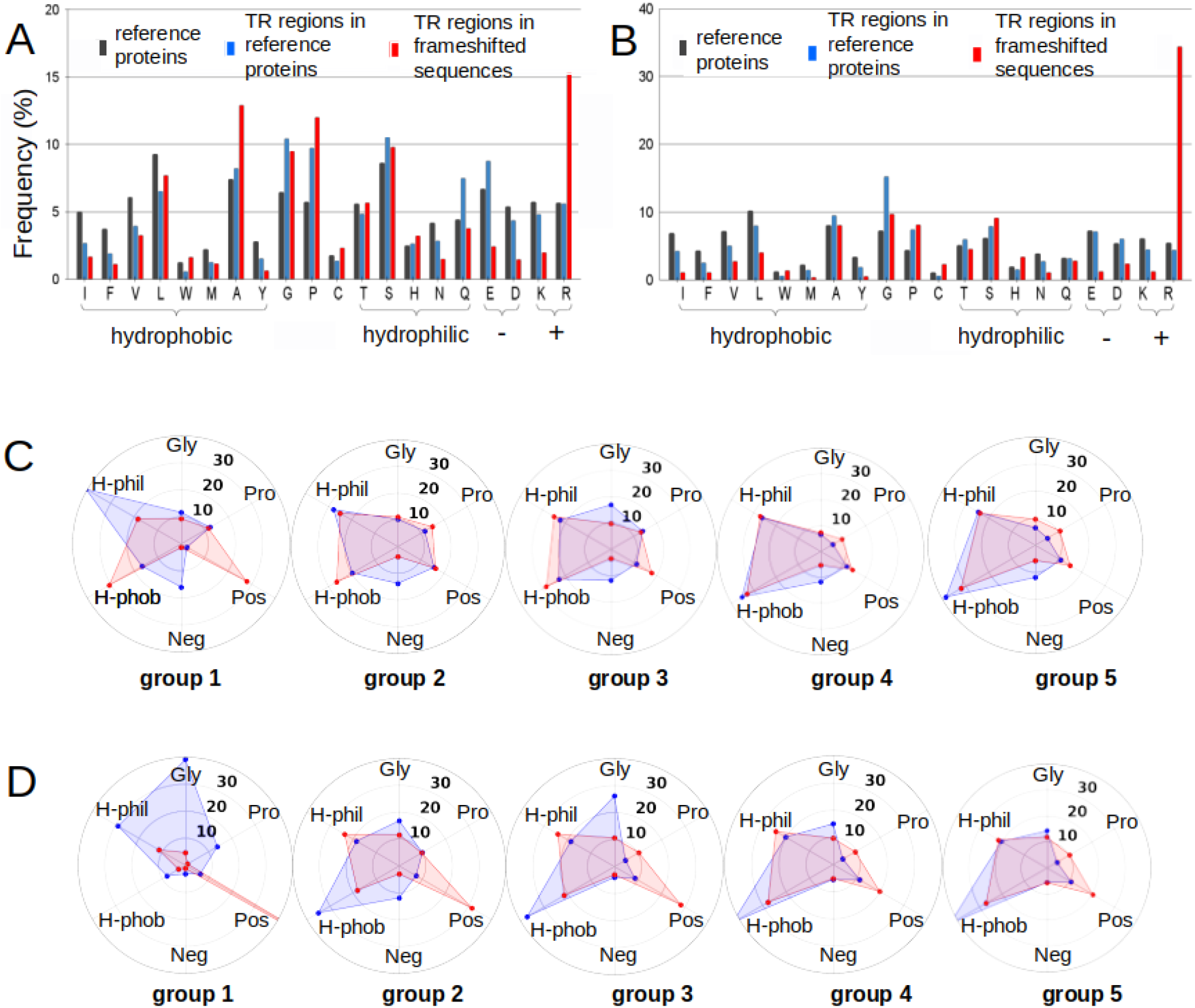
Histograms of AA composition of the reference and frameshifted sets of TRs in eukaryotes (A) and prokaryotes (B). AA frequencies (%) in all analyzed eukaryotes proteins (black bars), in TR regions of eukaryotic proteins (blue bars), and in frameshifted sequences of these TR regions (red bars). Hydrophobic residues are Ile, Phe, Val, Leu, Trp, Met, Ala, Tyr; hydrophilic residues are Ser, Thr, His, Asn, Gln; negatively charged are Glu, Asp; positively charged are Lys, Arg. Hereinafter, AA frequency of a specific AA = Total number of a specific AA in the set / Total number of all AAs in the set. Polygon graphs of AA composition (%) of the five TR groups in eukaryotes (C) and prokaryotes (D). H-Phob: hydrophobic, Gly: Glycine, Pro: Proline, H-Phil: hydrophilic, Neg: negatively charged, Pos: positively charged residues. Polygons of reference proteins and frameshifted sequences are in blue and red, respectively. For clarity of the graphs, the cysteine residue is not shown because it is rare in both sets (reference and frameshift).

In eukaryotes, the most drastic changes were observed in the occurrence of charged residues. In the frameshifted sequences, the frequency of the positively charged Arg substantially increases while the negatively charged residues decrease (Figure 2A). Another important observation was that the hydrophobicity of the frameshifted TR regions increases due to both the increase of apolar residues, especially Ala, and the decrease of polar residues, especially Gln. Thus, in the TR regions of eukaryotic proteins, the frameshift events, in general, lead to more hydrophobic and positively charged sequences.

In prokaryotes, the TR regions of the frameshifted sequences contain a considerably higher proportion of Arg compared to the TR regions in the reference proteins (Figure 2B). However, in contrast to eukaryotic organisms, in the frameshifted sequences of prokaryotes, the total hydrophobicity is reduced.

In summary, a prevailing trend in the frameshifted TRs is a notable increase in the positive charge when compared to their respective reference sequences in both prokaryotes and eukaryotes.

We also compared the AA composition of TR regions between the five groups described in the previous section. Our analysis demonstrated significant differences in the AA composition between reference and frameshifted sequences within each group (Figures 2C, 2D and Supplementary Tables S1 and S2).

*Group1*. In the reference proteins of eukaryotes, the homorepeats (HRs) are enriched (in order of increasing occurrence) in Gln, Ser, Glu, Pro, Ala, and Gly (Supplementary Table S1 and Figure S3). HRs are typically hydrophilic, with a higher prevalence of negative charges compared to positive ones (Figure 2C). In contrast, the frameshifted sequences are more hydrophobic, mostly due to the increase in Ala and Leu (Figure 2A). They also have a drastic drop in hydrophilic Gln and in negatively charged residues. The most remarkable increase was recorded for positively charged residues (Figure 2C), namely Arg (Figure 2A). In prokaryotes, this increase in Arg was observed, to an even greater extent (Figure 2D). Unlike eukaryotes, prokaryotic frameshifted HRs, similarly to the reference ones, have a very low percentage of hydrophobic residues.

At the DNA level, a specific amino acid HR is often encoded by the repetition of the same codon. Analyzing eukaryotic sequences, we observed the largest peak of codon usage at CAG (Gln). Eight other abundant codons have frequencies over 4%: GGC (Gly), GAG (Glu), TCC (Ser), GCC and GCG (Ala), CCG and CCA (Pro), and CAA (Gln) (Supplementary Figure S4). As a result of frameshifts, the most abundant codon CAG (Gln), is transformed into AGC (+1 shift) and GCA (-1 shift), corresponding to Ser and Ala, respectively - amino acids that are abundant in the frameshifted homorepeats (Supplementary Figures S5 and S6). The remarkable increase in poly-Arg within the frameshifted homorepeats, may be explained by the fact that Arg is encoded by six different codons, three of which, in the main frame, encode for Glu, Ser and Ala that are abundant in the reference homorepeats. The other AAs (Ser and Leu) encoded by six different codons in the frameshifted homorepeats, increase to a lesser extent due to the lower frequencies of the corresponding codons in the reference homorepeats (Supplementary Figures S4-S6).

*Group 2*. In eukaryotes, the 2-residue repeats of reference proteins are enriched in Ser, Pro, Arg, and Gly (Supplementary Table S1). Interestingly, among all TRs in the reference proteins, 2-residue repeats contain an unusually high percentage of Arg. Frameshifted TRs follow this trend, having Pro, Arg, Gly, and Ser as the most frequent residues. In general, AA compositions of reference and frameshifted sequences are similar. At the same time, unlike HRs, TRs from group 2 have almost the same percentages of positively and negatively charged residues in reference proteins. The frameshift leads to a drastic increase in positive charge of these TRs due to reduction of negatively charged residues, similar to those observed in group 1 (Figure 2C). The increase in hydrophobicity in the frameshifted TRs is also sustained, albeit to a lesser extent. In prokaryotes, the charged residues follow the same trend; however, the trend for hydrophobicity is reversed (Figure 2D).

*Group 3*. In TRs with 3-8 residue repeats, the tendencies associated with charged residues remain the same as in the groups 1 and 2, both in eukaryotes and prokaryotes (Figures 2C and 2D). The hydrophobicity of TRs in eukaryotic sequences is similar to their frameshifted sequences and even higher in the reference proteins of prokaryotes. In eukaryotes, the most frequent individual residue in the reference set is Gly, followed by Pro. This can be explained by the presence of collagen sequences in this group. Gly is also the most frequent in prokaryotes. The frameshifted sequences of both prokaryotes and eukaryotes are enriched in Arg.

*Group 4 and 5*. The TRs from these two groups have similar AA compositions in both eukaryotes and prokaryotes, Therefore, the conclusions mentioned below can be applied to both types of organisms. Unlike short repeats, those longer repeats exhibit increased hydrophobicity in reference proteins compared to the frameshifted sequences (Figures 2C and 2D). At the same time, the reference TRs continue to be predominantly composed of negatively charged residues, whereas the frameshifted TRs primarily contain positively charged ones. In general, the AA composition of the reference TRs becomes similar to the average composition of the analyzed proteomes (Figure 3A). In the frameshifted TRs, Arg continues to be the most common residue (Supplementary Tables S1 and S2).

**Figure 3.**
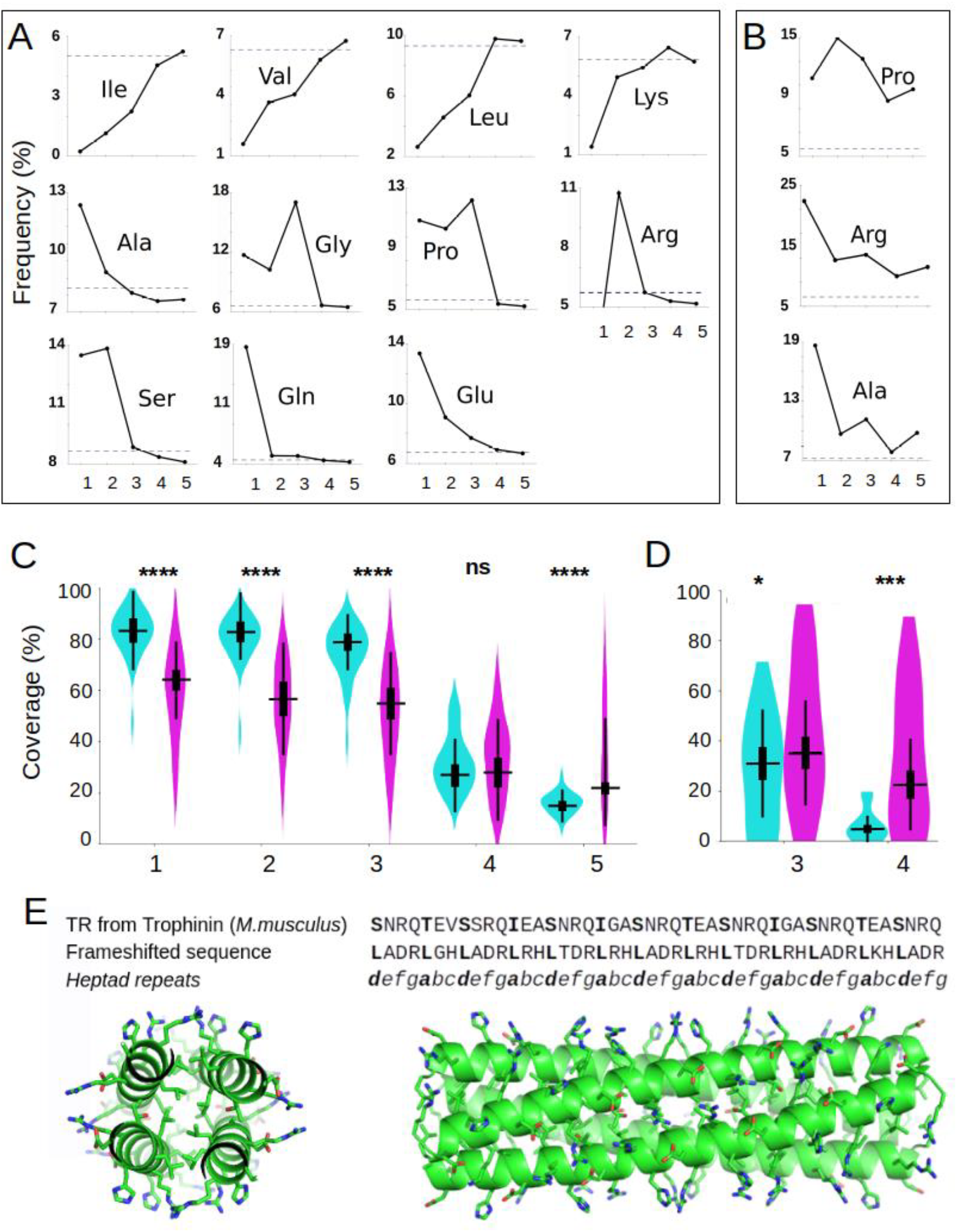
Line graphs of the most prominent changes in AA frequencies depending on the TR groups in eukaryotic proteins (A) in reference and (B) in their frameshifted sequences. Broken lines denote the average percentage of a given AA in reference proteomes. (C) Coverage of Intrinsically Disordered Regions (IDRs) within TR regions (percentage of AAs in TRs, which have unstructured conformation) of proteins in eukaryotic organisms and (D) in prokaryotes. The results for reference proteins and frameshifted sequences are shown in blue and magenta, correspondingly, and for each TR group separately. The observed variance is statistically significant between reference and frameshifted sets performed by the two-sided T-test. ****, *** and * mean p-value < 0.0001, p-value < 0.001 and p-value < 0.05, respectively. ns means non-significant. (E) Trophinin (Mus musculus) E9Q160_MOUSE. AlphaFold2 prediction of frameshifted sequence (pLDDT > 90). Axial and lateral projections of tetrameric structure.

Thus, generally, the frameshifted TRs from all the groups have more positively charged residues and less negatively charged residues compared to the reference TRs (Figures 2C and 2D). At the same time, the hydrophobicity of the short repeats increases in the frameshifted set and decreases with the growth of the repeat length. The frameshifted TRs are more hydrophobic than the reference ones due to the increase in Ala and to a lesser extent in Leu. They have a drastic drop in the frequency of polar Gln, negatively charged residues, and a strong prevalence of the positively charged Arg (Figures 3A and 3B, Supplementary Tables S1 and S2).

### 4. Typical repeats detected by clustering

We clustered the most occurring repeat motifs (see Methods). Clusters that comprise the greatest number of TRs sharing identical consensus sequences primarily consist of homorepeats (Supplementary Figure S7). In eukaryotes, as was expected, the top-ranked repeat motifs of the reference proteins obtained by the clustering correspond to the same most frequent AAs of the homorepeats revealed by AA composition analysis (Gln (7150 times), Gly (5020), Ala (4664), Pro (4649), Glu (3987) and Ser (3737)). The clustering of 2-residue repeats from group 2 showed that the most frequent pairs are Ser-Arg (2821), Pro-Ala (2358), Arg-Glu (2335), Ser-Pro (1727), Pro-Gly (1542), Gly-Ser (1487) and Glu-Lys (1427). This is generally in agreement with the AA composition of the 2-residue repeats, which are enriched in Ser, Pro, Arg, Gly, Glu, and Ala (Supplementary Table S1). At the same time, not all combinations of the most frequent AAs are present among the top-ranked repeats from group 2. For example, the negatively charged Glu-Ser, Glu-Gly, Glu-Ala, and Glu-Pro are not at the top of the list, in contrast to the positively charged TR formed by Ser-Arg and Arg-Gly, with Ser-Arg being the most frequent motif in group 2. This suggests that Arg-containing 2-residue repeats play an important functional role in the reference proteins. Group 3 exhibits an abundance of Gly and Arg, with the Gly-Pro-Pro motif representing the largest cluster (9487) not only within this group but also across all the groups (Supplementary Figure S7). This can be explained by the widespread occurrence of the collagen sequences in the reference proteins of eukaryotes. The clusters of TRs from group 4 and group 5 have significantly fewer members, and as a result, even the top-ranked TR sequences cannot be regarded as typical TR motifs.

For frameshifted eukaryotic sequences, the top-ranked repeat motifs of group 1 are homorepeats of Arg (6213), Ala (6174), Pro (2998), Ser (2726), and Gly (2255) (Supplementary Figure S7). The most significant distinction from the reference proteins is the emergence of positively charged Arg, coupled with the absence of negatively charged Glu and polar Gln. TRs from group 2 follow this trend, having apolar TRs with small side chains (Ala-Pro and Ala-Gly) and positively charged ones (Arg-Pro and Arg-Gly) as the most typical motifs. The TRs from group 3 are not among the 15 top-ranked motifs (Supplementary Figure S7). In comparison with the reference set, these TRs lack collagen-like motifs but primarily consist of positively charged residues and small-sized residues.

In prokaryotes, TR sets are significantly smaller. Within the reference TRs, homorepeats from group 1 are less common, with only three of them (Gly, Gln, and Pro) appearing in the 15 top-ranked motifs (Supplementary Figure S7). In group 2, the most prevalent motifs consist of combinations of small-sized residues along with negatively charged ones, such as Asp-Leu, Pro-Ala, Gly-Ser, and Glu-Pro. Surprisingly, one of the most frequent motifs belongs to group 3. These motifs (Gly-Gly-Ala, Gly-Gly-Asn, Gly-Gly-Thr) are commonly observed in bacterial cell surface proteins from the PE family (39). The typical TR motifs of the frameshifted sequences significantly differ from the motifs of the reference proteins, with a pronounced prevalence of Arg homorepeats (Supplementary Figure S7). Additionally, other frequently occurring motifs belong to group 2 and group 3 (such as Arg-Gly, Arg-Pro, Arg-Ser, Arg-Arg-Gly, and Arg-His), all of which are also notably enriched in Arg.

Thus, despite variation in TR motifs between prokaryotes and eukaryotes in the reference set, our clustering revealed in frameshifted sequences remarkable dominance of poly-Arg, Pro-Arg and Gly-Arg, regardless of the species, clearly indicating that the frameshifts favor the enrichment of positively charged Arg.

### 5. Intrinsic Disorder Within TR Regions

Intrinsically Disordered Regions (IDRs) are important attributes that enable assessment of the structural states and functional roles of the analyzed proteins. We predicted IDRs by using IUPred (30) and observed that the coverage of IDRs in the reference TR regions from eukaryotes and prokaryotes sets are 57.20% and 21.24 %, respectively. Notably, these percentages decrease to 47.26% in eukaryotes and increase to 31.86% in prokaryotes in the frameshifted sets. We observed that the shorter the repeats, the higher the disorder (Figures 3C and 3D). For eukaryotes, the highest coverage of IDRs was observed in homorepeats from the reference set. The TRs from groups 2 and 3 (up to 8 residue-long repeats) have a similarly high level of IDRs (Figure 3C). The coverage of IDRs drops dramatically within TRs with repeats of more than 8 residues (groups 4 and 5). TRs of the frameshifted sequences from eukaryotes are less disordered than the reference ones in groups 1, 2, and 3. In groups 4 and 5, while the IDR coverage of the frameshifted TRs decreases in comparison with groups 1-3, it becomes slightly higher than that of the reference set.

In our prokaryotic set, the number of IDRs in the TR is insufficient to draw statistically valid conclusions for group 1, 2 and 5. Therefore, Figure 3D shows the results from group 3 and 4 only, which had a sufficient number of IDRs to get statistically significant differences. In group 3, we noticed a slight increase in disorder within the frameshifted sets when compared to the reference ones, which stands in contrast to the trend observed in eukaryotes. The higher disorder in the frameshifted set than in the reference set is even more pronounced in group 4. Similar difference was observed in group 4 of eukaryotic species.

### 6. In search of the 3D structures predicted by AlphaFold2 among frameshifted sequences

The majority of the frameshifted sequences are predicted to be intrinsically disordered (17). In agreement with this result, we also predicted a large number of IDRs in the frameshifted TR sequences, particularly within groups 4 and 5 (Figure 3C). Still, some of the sequences were predicted to be structured, prompting our further exploration to understand the atomic details of the formed 3D structures. For this purpose, we used the AlphaFold2 tool (see Materials and Methods). In our previous analysis of protein isoforms, AF2 modeling suggested that a frameshift in one out of several exons of a gene frequently does not disrupt the overall protein structure (40). Here, we analyzed the entire frameshifted TR regions in search of 3D structural models with high confidence scores. We selected a non-redundant set of 14 031 TR-containing frameshifted sequences, which were found at least two times across 29 eukaryotic species (see Materials and Methods). Therefore, we called them “conserved”. Moreover, these sequences were predicted by TAPASS (27) to be structured. This set was used as input for our large-scale AF2 modeling.

Among obtained AF2 structural models of frameshifted TR regions, only 459 had a high confidence score pLDDT ≥ 70. They were analyzed manually by PyMol (42). A few of them, mostly Zn-finger and ankyrin containing sequences, had a high confidence score of AF2 in the frameshifted sequences, while the corresponding reference sequences were predicted to be IDRs. It turned out that these reference proteins have low-quality annotation in the databases, which was not supported by RNAseq or mass-spectral data. This suggested that the reference sequences may be erroneously annotated and, in fact, the frameshifted ones should be in the reference set. Still, some of the frameshifted structures were selected for subsequent experimental evaluation of predicted structures. For example, several frameshifted high-quality annotated sequences have well-distinguished collagen-like and α-helical coiled coil TRs. The corresponding reference sequences of most of them are predicted to have IDRs. However, in certain cases, such as in Trophinin (*Mus musculus*) E9Q160_MOUSE (Figure 3E), AF2 models represent α-helical coiled coil oligomers with pLDDT > 90 in both reference sequence and its frameshifted variant. This suggests that in some cases of short repeats, a frameshift event can give rise to novel structures and functions.

### 7. Propensity of TR regions to aggregate

Some proteins tend to aggregate and, upon changes in the conditions, form insoluble deposits. In certain cases, these aggregates play functional roles (42), and in others they are linked to amyloidosis or other diseases (43-44). TR regions of several reference proteins, such as Huntingtin, α-synuclein, and yeast prions, form aggregates. It was interesting to test how the frameshift in TRs changes their aggregation potential. For this purpose, we predicted the aggregation potential of the reference and frameshifted TRs. TANGO and ArchCandy predictors (28-29) were selected based on their performance and the fundamental differences in their algorithms. We focused on the eukaryotic sets because the limited number of TRs with aggregation-prone regions (ARs) in prokaryotic sequences renders them unsuitable for statistical tests.

TANGO predicted a higher aggregation propensity in eukaryotic frameshifted TRs compared to the reference ones for repeats shorter than 9 residues (Figure 4A), while the opposite trend was observed for longer repeats. ArchCandy predicted similar AR coverage for both reference and frameshifted TRs from groups 1 to 3, showing a steady decrease in ARs with the increase in repeat length (Figure 4B). The situation changes for TR with repeats of more than 8 residues (group 4 and 5), where the reference TRs become more aggregation-prone than the frameshifted ones.

**Figure 4.**
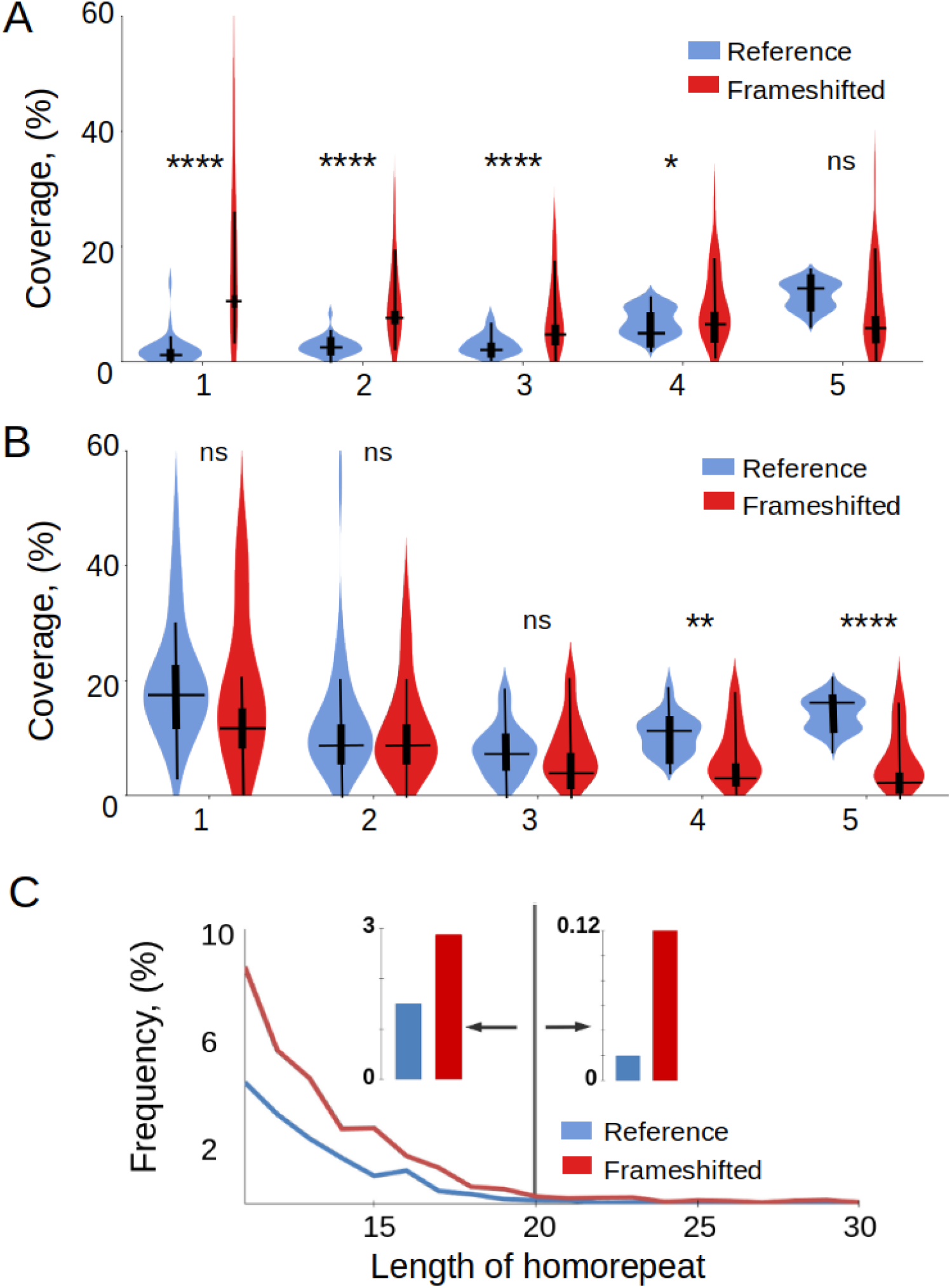
Coverage of aggregation-prone regions (ARs) within TR regions of the proteins in eukaryotic organisms predicted by TANGO (A) and ArchCandy (B) Results for reference sequences are in blue and for frameshifted ones in red. The coverage of ARs in TR regions was calculated by summing up all predicted AR residues overlapping with TR regions divided by the total number of AA in the TR regions of the set. The observed variance performed by the two-sided T-test is statistically significant in TR groups, except group 5. **** and * mean p-value < 0.0001 and p-value < 0.05, respectively. ns means non-significant. (C) Length distribution of hydrophobic homorepeats (I, F, V, L, W, M, A, Y) in reference and frameshifted sets of eukaryotes. The values on the graph are calculated as the number of hydrophobic homorepeats of a certain length divided by the total number of all detected homorepeats. The vertical line subdivides the homorepeats into those that have high aggregation potential (lengths from 11 to 20 residues) and homorepeats able to form transmembrane helices (from 21 and more). The insets on both sides from the vertical line show the difference between the average frequencies of the reference and frameshifted homorepeats within these two length ranges.

Thus, both predictors agreed that TRs with long repeats from group 5 are more aggregation-prone in the reference than in the frameshifted sequences. This result can be attributed to the higher hydrophobicity of the reference TRs compared to the frameshifted set. The results for the short repeats are mixed and depend on the predictor. This can be explained by the algorithm differences: TANGO favors hydrophobic residues within ARs, whereas ArchCandy, in addition to hydrophobic residues, takes into account high aggregation potential of Gln and Asn-rich regions. Poly-Gln is one of the most frequent reference homorepeats in eukaryotes (19) (Figure 3A and Supplementary Table S1). This can explain ArchCandy prediction, indicating that the aggregation of homorepeats from the reference set is as high as in the frameshifted set. The significance of Gln/Asn for the aggregation of short repeats became even more evident when we predicted exposed aggregation-prone regions (EARs) located in disordered regions of proteins and accessible for intermolecular interactions (27). ArchCandy predicted that EARs are more frequent in the reference than in the frameshifted TRs of short repeats (Supplementary Figures S8 and S9). As the Gln frequency in the reference TRs of groups 2 and 3 decreases, the aggregation potential predicted by ArchCandy also decreases. TANGO cannot detect aggregation of Gln- or Asn-rich regions and, therefore, predicts very small numbers of EARs in both reference and frameshifted sequences.

Merely counting ARs does not provide a realistic answer to the question of which sequences, whether reference or frameshifted, are more prone to aggregation. This is especially true for TRs from groups 4 and 5. Indeed, the structural context of proteins plays a very important role in the aggregation potential of sequence motifs (27). Frequently, ARs with a high proportion of hydrophobic residues are involved in the stable 3D structures of proteins. These ARs can be hidden in the structure and inaccessible for intermolecular aggregation. This is particularly true for the reference sequences that have been evolutionarily selected to form specific stable functional structures. Importantly, the principle of evolutionary selected sequence does not apply to the majority of the frameshifted sequences. Even with an amino acid composition that is hydrophobic enough to fold into a stable 3D structure, these sequences were not evolutionarily shaped for this folding. Our large-scale AF2 modeling of TR from the reference and frameshifted sequences supported this conclusion, as we did not find many frameshifted sequences predicted to fold in a plausible 3D structure (see previous section). Thus, while ArchCandy and TANGO predicted a slightly higher aggregation potential for reference sequences in TR groups 4 and 5 compared to frameshift sequences, in reality, the latter may be more prone to misfolding, thus contributing to their preferential aggregation.

### 8. Implications of hydrophobic homorepeat relocalization during frameshifting events

If homorepeats consist of hydrophobic AA, they tend to aggregate. However, when the length of a homorepeat exceeds 20 residues, it can be inserted into the membrane, form transmembrane (TM) α-helices and avoid aggregation (27). We analyzed the frequency of hydrophobic homorepeats depending on their length and found that frameshifted sequences have more hydrophobic homorepeats than reference sequences across all length ranges (Figure 4C). This is especially true for homorepeats exceeding 20 residues with potential to form TM helices. We observed that often during a frameshift, either hydrophobic homorepeats in the reference sequence turn into hydrophilic ones, or, conversely, hydrophilic ones turn into hydrophobic ones. Consequently, frameshifting in the homorepeat region can alter the solubility or insolubility of certain proteins and (or) their localization, whether within the membrane or elsewhere. Given that in general the frameshift increases the number of hydrophobic homorepeats, it should lead to the increase in aggregation and location in the membranes.

## 9. DISCUSSION AND CONCLUSIONS

Our large-scale comparative analysis of reference and frameshifted sequences within TRs revealed a number of intriguing correlations. For example, we observed that the disparity in physicochemical characteristics between reference and frameshifted TR sequences is greatly influenced by the length of the repeats. Smaller repeats exhibit more pronounced differences, whereas repeats with a length of 50 residues or more tend to converge with their characteristics resembling both each other and the average values found across the entire proteome.

In the case of short repeats, we observed that frameshifted sequences exhibit greater hydrophobicity and a reduced number of IDRs. What makes this result particularly interesting is the fact that, in general, the frameshift sequences have more IDRs than the reference ones (17). In addition, our analysis of hydrophobic homorepeats based on their length revealed that the frameshifted sequences contain a higher number of predicted aggregation-prone regions and transmembrane helices compared to the reference sequences. In numerous instances, hydrophobic homorepeats are absent in the reference set but appear in the frameshifted set, or vice versa. As a result, frameshifted homorepeats can render certain proteins insoluble or impart membrane localization to them.

In regard to changes in AA composition, the most prominent difference is a substantially higher percentage of Arg in the frameshifted TRs than in the reference ones being approximately three times greater in eukaryotes and about seven times higher in prokaryotes. Among the most frequent repetitive motifs of the frameshifted sequences are poly-Arg and di-peptide repeats Gly-Arg and Pro-Arg. In the reference proteins, Arg-rich repeats play various functional roles. For example, Arg-rich Intracellular Delivery (AID) peptides transport big molecules into cells in plants using both covalent and noncovalent protein transductions (45). The C-terminal part of U1-70K protein, a small nuclear ribonucleoprotein particle, which contains Arg-Ser dipeptides region, may be involved in the regulation of mRNA splicing in Drosophila and pre-mRNA splicing in plants (46-47). Mammalian heterogeneous nuclear ribonucleoprotein (hnRNP) E1B-AP5 is methylated *in vivo* within Arg-Gly-Gly (RGG)-box motifs. E1B-AP5 is recognized for its role in mediating protein-RNA interactions (48). It has been also found that Arg-rich proteins can form liquid-like droplets through Liquid-Liquid Phase Separation (LLPS). Arginine can switch between dense and liquid phases at high salt concentrations, but not lysine (49). Thus, arginine may be used to control droplet behavior (49). We found that Arg-containing 2-residue TRs are especially frequent in the reference proteins. A number of studies suggest that these TRs have specific functions being involved in protein binding to DNA and RNA, participating in the organization of an extracellular matrix, and being important elements of the LLPS process (50). For example, Arg-Ser di-peptide repeats are involved in alternative splicing control, Arg-Gly in transcriptional regulation RNA binding, Arg-Val regions in gene carriers, and Arg-Glu in cell survival signaling (51). The functional significance of these regions in the reference proteins suggests that frameshifted Arg-rich TRs, once they arise, may confer new functions to the protein.

An imbalanced increase of Arg-rich TR in frameshifted sequences may also have a significant pathological impact if they are expressed in the organisms. Indeed, it has been shown that Arg-rich sequences are highly toxic in cell and animal models of Amyotrophic lateral sclerosis (ALS) (52-54). The same di-peptides may affect ribosome-associated quality control (55). Furthermore, Pro- and Arg-rich peptides have been demonstrated as allosteric inhibitors of 20S proteasome (56). Arg-rich peptides are also modulators of protein aggregation and cytotoxicity playing a role in Alzheimer’s Disease (57). In addition, *de novo* frameshifts in HMGB1 protein, which replace the intrinsically disordered acidic tail of HMGB1 with an Arg-rich basic tail, cause brachyphalangy, polydactyly, and tibial aplasia/hypoplasia syndrome (BPTAS) (58). Thus, our finding of the exceptionally high abundance of Arg as a result of frameshifting in TRs points out to a potential link of the frameshifts with pathological effects.

Other common di-peptide repeats identified in frameshifted sequences, lacking Arg, are also known to play important functional roles in reference proteins. For example, Gly-Ser repeats in Chimeric antigen receptor (CAR) linkers provide the flexibility necessary for antigen-binding sites to change conformation (59).

We compared the aggregation tendencies of frameshifted sequences with short repeats (groups 1, 2 and 3) to those of the reference set. Quantitatively, their aggregation tendencies were similar; however, qualitatively, the high aggregation potential of frameshifted sequences is due to a high content of hydrophobic residues, while the high aggregation potential of the reference sequences is due to abundant poly-Q and poly-N repetitive sequences.

Previously, we found protein isoforms with frameshifts within a single repeat in a TR structural domain. AF2 modeling suggested that these single-repeat frameshifts do not disrupt the overall structure of the domains composed of the TRs. The frameshifted repeat can fit into the remaining structure (40). However, in the present study, when we sought AF2 structural models with high confidence for cases where the frameshift covers the entire TR regions, we obtained ambiguous results. The vast majority of the models had low confidence scores. This was especially true for the long repeats belonging to Class III, IV, and V of the structural classification (37). Among the TR regions formed by short repeats from Class II, we found several AF2 models with high confidence scores featuring either a collagen triple helix or α-helical coiled-coil oligomers. A subset of the most interesting frameshifted structural models was chosen for subsequent experimental evaluation, and the results will be reported upon completion.

Thus, our analysis revealed that frameshift can have distinct and significantly stronger effects on TR regions compared to non-repetitive sequences. While previous studies have indicated that, despite frameshift causing differences in protein sequences, several of their characteristics, including their hydrophobicity profiles and intrinsic disorder profiles, maintain similarity in the corresponding frameshifted sequences (17), our current study demonstrates that frameshifts in TR regions, particularly those composed of short repeats, lead to drastic changes. These changes include increased hydrophobicity, Arg-rich sequences, novel aggregation-prone regions, and transmembrane helices. Consequently, frameshifts in TRs may be most crucial for altering binding partners, cellular localization, and other functional roles. Moreover, these frameshifts can pose significant risks and contribute to various pathologies.

## ACKNOWLEDGMENTS

We thank Drs Layla Hirsh Martínez, Damien Devos, Peter Tompa, Nikola Arsic for discussion, careful reading of the manuscript, and valuable comments. The AF2 modeling was done by using HPC resources from GENCI-IDRIS (Grant 2023-AD010313976R1) supported by the Grand Equipement National de Calcul Intensif (GENCI) and the Centre National de la Recherche Scientifique (CNRS), under the aegis of the French Ministry of Higher Education and Research.

