## Supplemental Data for "Frameshifts in Tandem Repeats: Consequences on Protein Physicochemical Properties and Function"

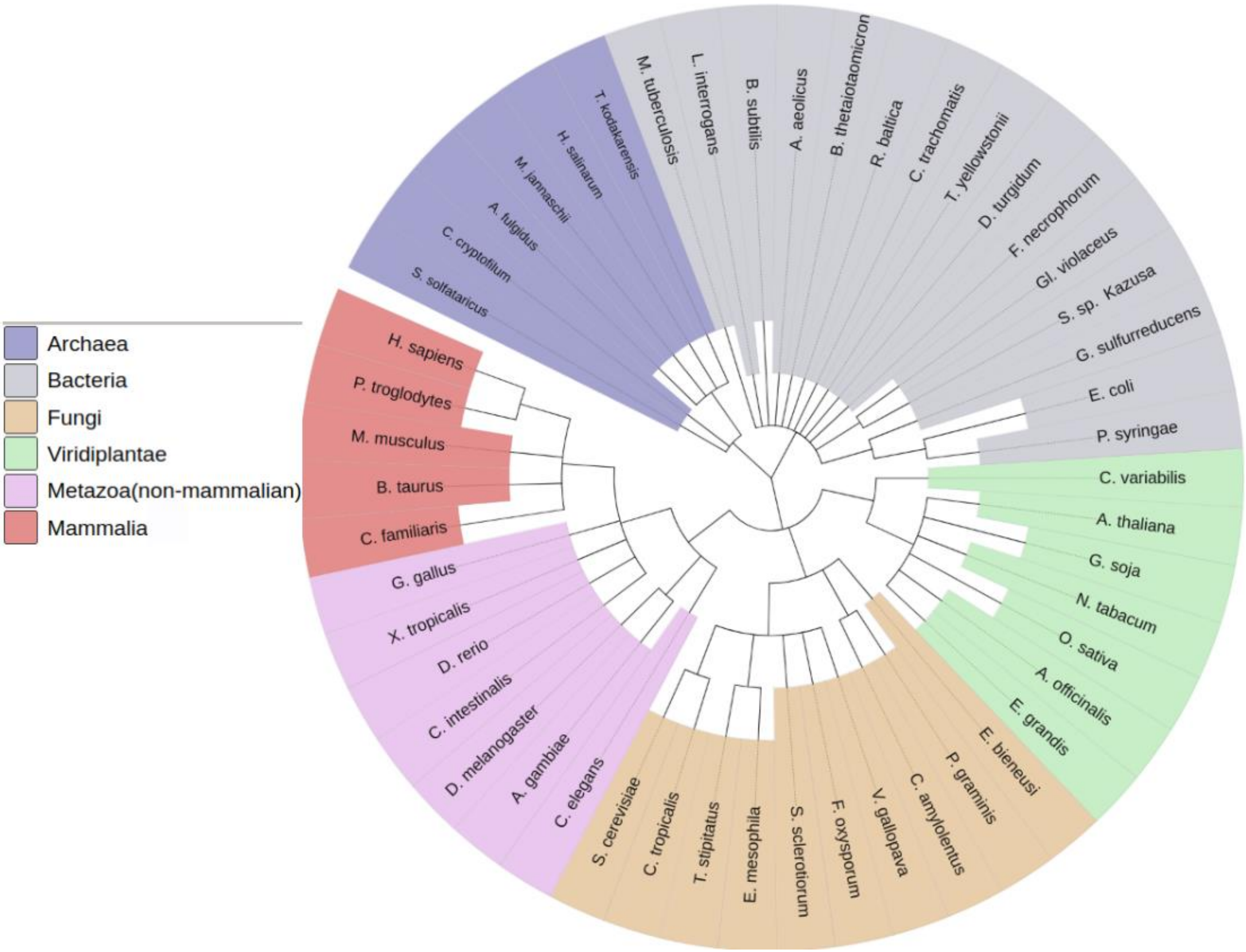


**Figure S1.** Phylogenetic tree of 50 species that were selected for the analysis. The tree is generated by iTOL.

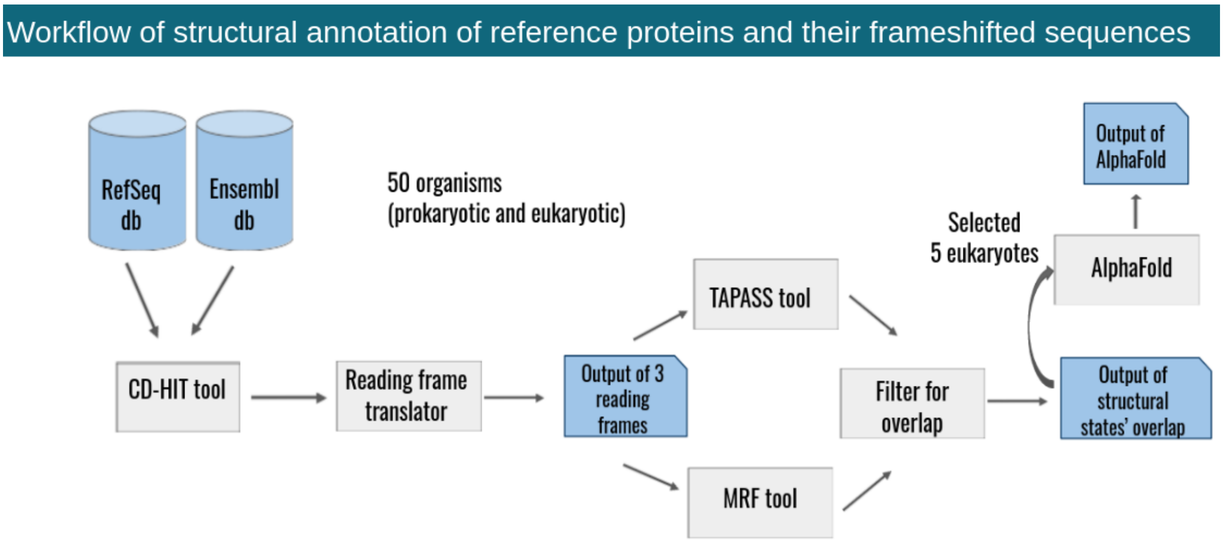


**Figure S2.** Workflow for annotation of reference proteins and their frameshifted sequences. CDSs of 50 prokaryotic and eukaryotic species obtained from RefSeq and Ensembl. The CD-HIT tool was used for clustering in order to remove duplication of sequences retrieved from RefSeq and Ensembl databases. Then, the non-redundant CDSs were processed by the Reading frame translator tool to get AA sequences in 3 reading frames longer than 60 AA. Here 3 reading frames are the main reading frame called reference frame, -1 and +1 shifted frames. Afterward, TRs were detected by MRF and the structural states of protein sequences were predicted by TAPASS. The result files are processed by an in-house script named Filter for overlap to compare the structural states of reference proteins and their frameshifted sequences. Finally, AF2 structural models were predicted for selected frameshifted sequence sets containing TR-containing proteins from five species.

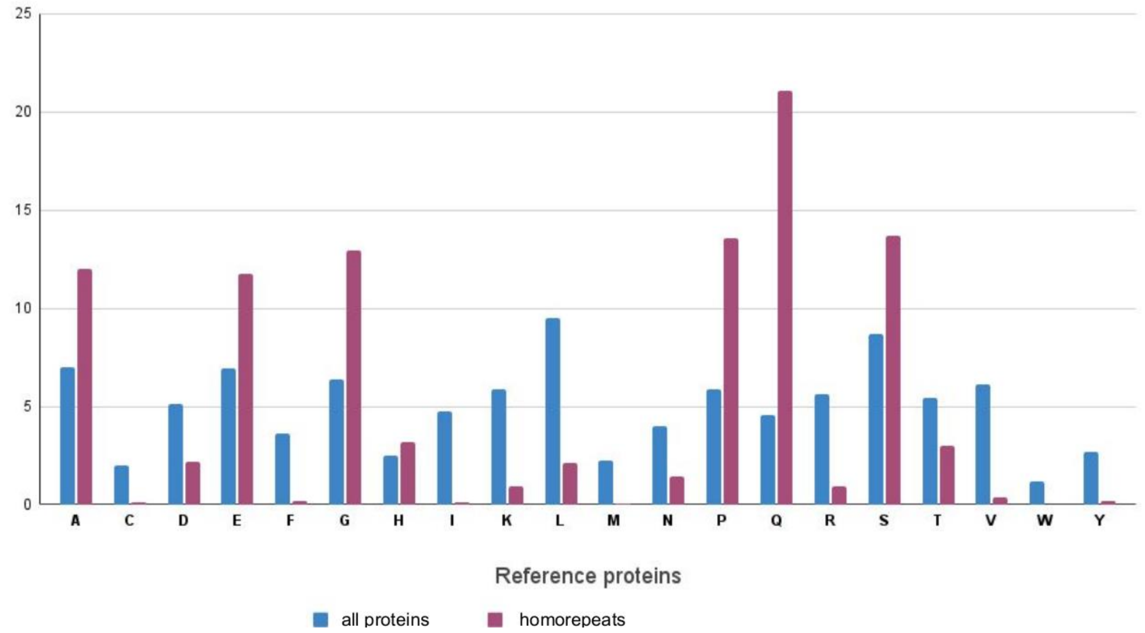


**Figure S3.** Frequencies of AAs (in %) in all reference proteins (non-repetitive and repetitive) (in blue) and in homorepeats (in red).

**Table S1.** Coverage (%) of AAs in TR groups in reference and frameshifted sequences of eukaryotes

| | Coverage of amino acids in tandem repeat groups | | | | | | | | | | | |
|---|---|---|---|---|---|---|---|---|---|---|---|---|
| | Eukaryotes reference sequences | | | | | | Eukaryotes frameshifted sequences | | | | | |
| Amino acid | group 1 | group 2 | group 3 | group 4 | group 5 | mean of groups | group 1 | group 2 | group 3 | group 4 | group 5 | mean of groups |
| Ala | 12.26 | 8.34 | 7.13 | 6.64 | 6.74 | 8.22 | 22.54 | 10.59 | 12.55 | 8.14 | 10.74 | 12.91 |
| Cys | 0.18 | 0.85 | 0.78 | 2.83 | 2.23 | 1.37 | 0.75 | 2.33 | 2.23 | 3.31 | 3.08 | 2.34 |
| Asp | 2.72 | 4.75 | 4.14 | 4.86 | 5.31 | 4.36 | 0.28 | 1.00 | 1.30 | 2.16 | 2.57 | 1.46 |
| Glu | 13.35 | 9.09 | 7.71 | 6.93 | 6.68 | 8.75 | 1.08 | 2.72 | 2.15 | 3.13 | 3.09 | 2.43 |
| Phe | 0.29 | 0.79 | 1.32 | 3.33 | 3.88 | 1.92 | 0.45 | 0.76 | 0.82 | 2.09 | 1.41 | 1.10 |
| Gly | 11.72 | 10.25 | 17.02 | 6.66 | 6.48 | 10.43 | 9.34 | 11.20 | 9.85 | 7.40 | 9.72 | 9.50 |
| His | 3.35 | 2.68 | 1.49 | 3.19 | 2.47 | 2.63 | 1.23 | 3.46 | 2.80 | 4.11 | 4.53 | 3.23 |
| Ile | 0.23 | 1.13 | 2.23 | 4.54 | 5.23 | 2.67 | 0.69 | 1.57 | 1.36 | 2.90 | 1.84 | 1.67 |
| Lys | 1.41 | 4.95 | 5.44 | 6.45 | 5.74 | 4.80 | 1.12 | 1.79 | 2.14 | 2.97 | 1.95 | 1.99 |
| Leu | 2.64 | 4.58 | 6.03 | 9.76 | 9.61 | 6.52 | 5.01 | 7.72 | 7.20 | 9.61 | 8.94 | 7.70 |
| Met | 0.23 | 0.75 | 1.26 | 1.92 | 2.06 | 1.24 | 0.60 | 0.92 | 1.17 | 1.82 | 1.27 | 1.16 |
| Asn | 1.65 | 2.01 | 1.97 | 4.47 | 4.13 | 2.85 | 0.87 | 1.38 | 1.49 | 2.32 | 1.36 | 1.48 |
| Pro | 12.34 | 11.66 | 14.00 | 5.46 | 5.26 | 9.74 | 11.58 | 14.95 | 13.20 | 9.67 | 10.64 | 12.01 |
| Gln | 18.68 | 5.04 | 5.02 | 4.47 | 4.26 | 7.49 | 2.28 | 3.49 | 3.44 | 4.78 | 4.83 | 3.76 |
| Arg | 1.06 | 10.75 | 5.75 | 5.31 | 5.18 | 5.61 | 26.72 | 14.55 | 15.68 | 11.23 | 13.12 | 16.26 |
| Ser | 13.48 | 13.83 | 8.85 | 8.35 | 8.10 | 10.52 | 10.06 | 9.69 | 11.24 | 9.84 | 8.08 | 9.78 |
| Thr | 3.43 | 4.35 | 5.05 | 5.73 | 5.68 | 4.85 | 4.08 | 7.00 | 6.05 | 6.15 | 5.07 | 5.67 |
| Val | 0.70 | 3.11 | 3.58 | 5.58 | 6.68 | 3.93 | 0.88 | 2.95 | 2.94 | 4.75 | 4.83 | 3.27 |
| Trp | 0.05 | 0.27 | 0.22 | 0.90 | 1.33 | 0.55 | 0.32 | 1.42 | 1.80 | 2.47 | 2.13 | 1.63 |
| Tyr | 0.26 | 0.84 | 1.03 | 2.61 | 2.95 | 1.54 | 0.15 | 0.54 | 0.60 | 1.16 | 0.81 | 0.65 |

**Table S2.** Coverage (%) of AAs in TR groups in reference and frameshifted sequences of prokaryotes

| | Coverage of amino acids in tandem repeat groups | | | | | | | | | | | |
|---|---|---|---|---|---|---|---|---|---|---|---|---|
| | Prokaryotes reference sequences | | | | | | Prokaryotes frameshifted sequences | | | | | |
| Amino acid | group 1 | group 2 | group 3 | group 4 | group 5 | mean of groups | group 1 | group 2 | group 3 | group 4 | group 5 | mean of groups |
| Ala | 5.74 | 11.17 | 13.42 | 8.49 | 8.67 | 9.50 | 1.39 | 7.93 | 11.06 | 9.37 | 10.55 | 8.06 |
| Cys | 0.23 | 0.21 | 0.48 | 1.01 | 0.95 | 0.58 | 0.37 | 2.38 | 2.16 | 3.68 | 3.01 | 2.32 |
| Asp | 3.28 | 11.97 | 4.03 | 5.52 | 5.56 | 6.07 | 0.74 | 2.11 | 1.91 | 3.20 | 3.90 | 2.37 |
| Glu | 8.56 | 6.21 | 6.86 | 7.31 | 6.68 | 7.12 | 0.46 | 0.86 | 1.10 | 1.98 | 1.86 | 1.25 |
| Phe | 0.23 | 1.88 | 2.25 | 4.07 | 4.22 | 2.53 | 0.00 | 0.79 | 0.85 | 2.18 | 1.68 | 1.10 |
| Gly | 30.60 | 10.71 | 19.37 | 7.75 | 7.72 | 15.23 | 4.73 | 11.62 | 10.66 | 9.71 | 11.97 | 9.73 |
| His | 1.17 | 1.72 | 0.85 | 1.82 | 1.97 | 1.51 | 0.74 | 3.73 | 3.46 | 3.98 | 5.08 | 3.40 |
| Ile | 0.35 | 3.64 | 4.21 | 6.70 | 6.55 | 4.29 | 0.09 | 1.05 | 0.91 | 2.09 | 1.50 | 1.13 |
| Lys | 2.93 | 3.86 | 4.63 | 5.82 | 5.18 | 4.48 | 0.46 | 0.87 | 1.34 | 2.17 | 1.25 | 1.22 |
| Leu | 0.47 | 10.12 | 8.87 | 10.23 | 10.18 | 7.97 | 0.46 | 3.85 | 3.64 | 5.92 | 6.25 | 4.03 |
| Met | 0.35 | 1.62 | 1.27 | 1.93 | 2.06 | 1.45 | 0.00 | 0.49 | 0.43 | 0.78 | 0.42 | 0.42 |
| Asn | 0.70 | 1.88 | 3.42 | 4.01 | 3.79 | 2.76 | 0.09 | 1.16 | 1.17 | 1.88 | 1.31 | 1.12 |
| Pro | 13.83 | 10.02 | 4.69 | 4.09 | 4.47 | 7.42 | 0.93 | 9.75 | 10.41 | 9.47 | 10.07 | 8.12 |
| Gln | 3.75 | 2.38 | 3.24 | 3.35 | 3.35 | 3.21 | 1.02 | 2.65 | 2.82 | 3.64 | 3.97 | 2.82 |
| Arg | 3.52 | 3.45 | 4.06 | 5.44 | 5.47 | 4.39 | 77.94 | 30.61 | 26.91 | 17.68 | 19.15 | 34.46 |
| Ser | 15.36 | 6.05 | 6.09 | 6.04 | 6.03 | 7.91 | 9.27 | 10.19 | 9.96 | 9.39 | 7.09 | 9.18 |
| Thr | 8.09 | 6.50 | 5.04 | 5.02 | 5.30 | 5.99 | 0.28 | 6.01 | 6.71 | 5.58 | 4.23 | 4.56 |
| Val | 0.23 | 5.14 | 5.53 | 7.15 | 7.29 | 5.07 | 0.56 | 2.35 | 2.76 | 3.96 | 4.32 | 2.79 |
| Trp | 0.12 | 0.33 | 0.44 | 1.07 | 1.25 | 0.64 | 0.37 | 1.27 | 1.32 | 2.34 | 1.59 | 1.38 |
| Tyr | 0.47 | 1.14 | 1.24 | 3.20 | 3.31 | 1.87 | 0.09 | 0.35 | 0.45 | 1.03 | 0.81 | 0.54 |

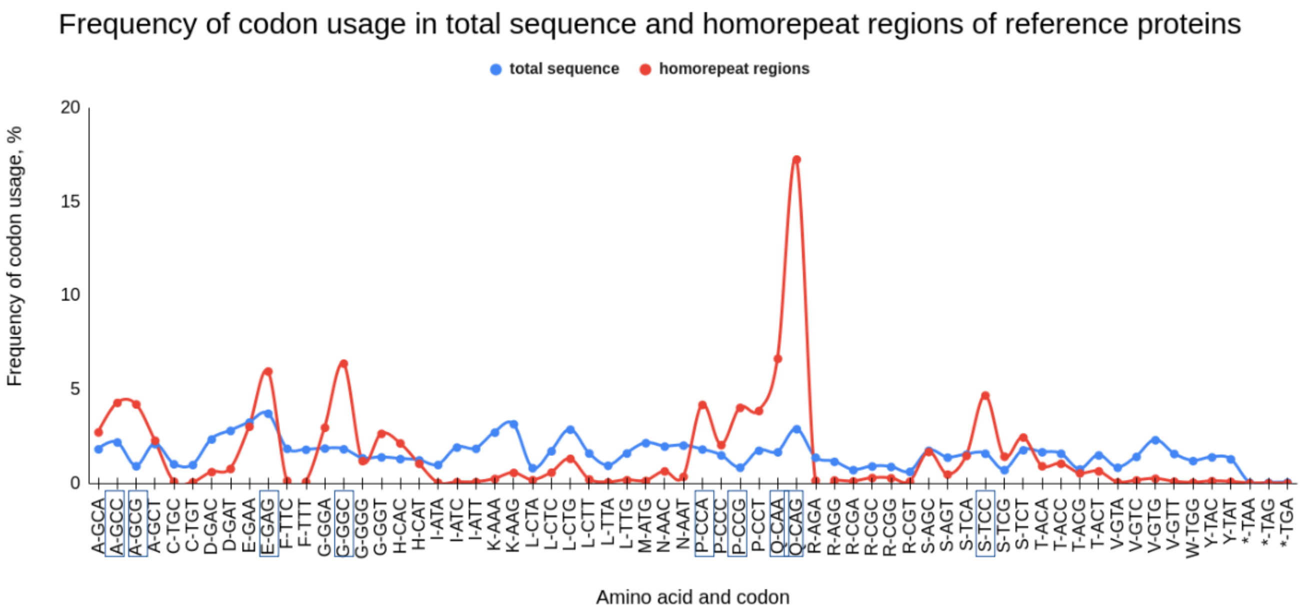


**Figure S4**. Codon usage in all reference proteins (blue) and homorepeats (red). The most frequent codons in homorepeats are marked by rectangles. The largest peak is at CAG codon (poly-Gln).

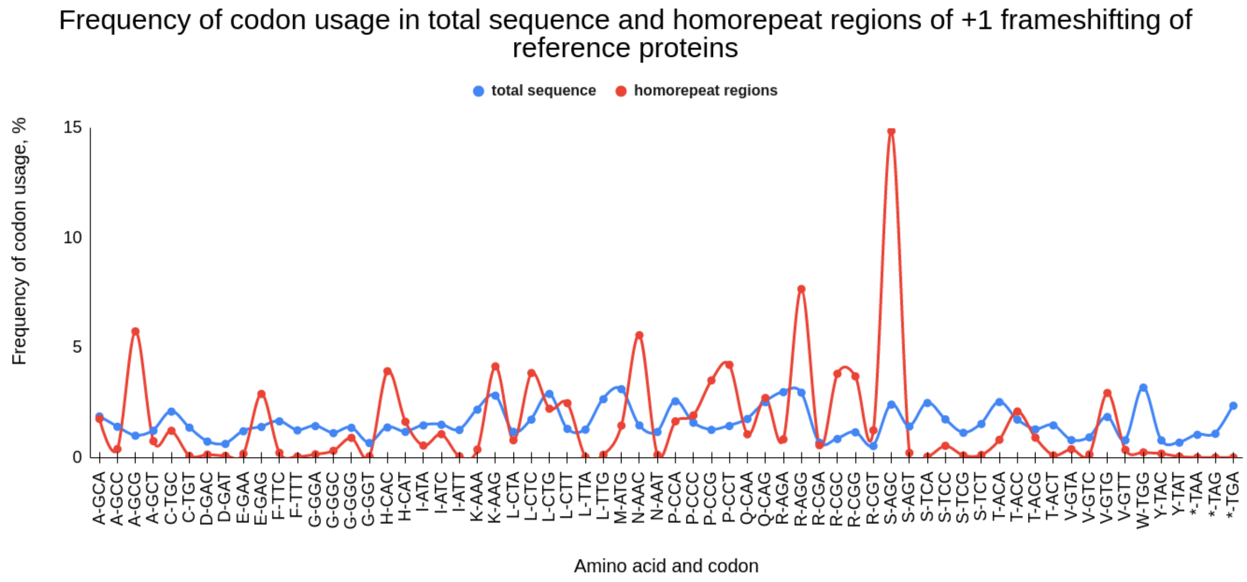


**Figure S5**. Codon usage in all +1 shifted reference proteins (blue) and +1 shifted homorepeats (red).

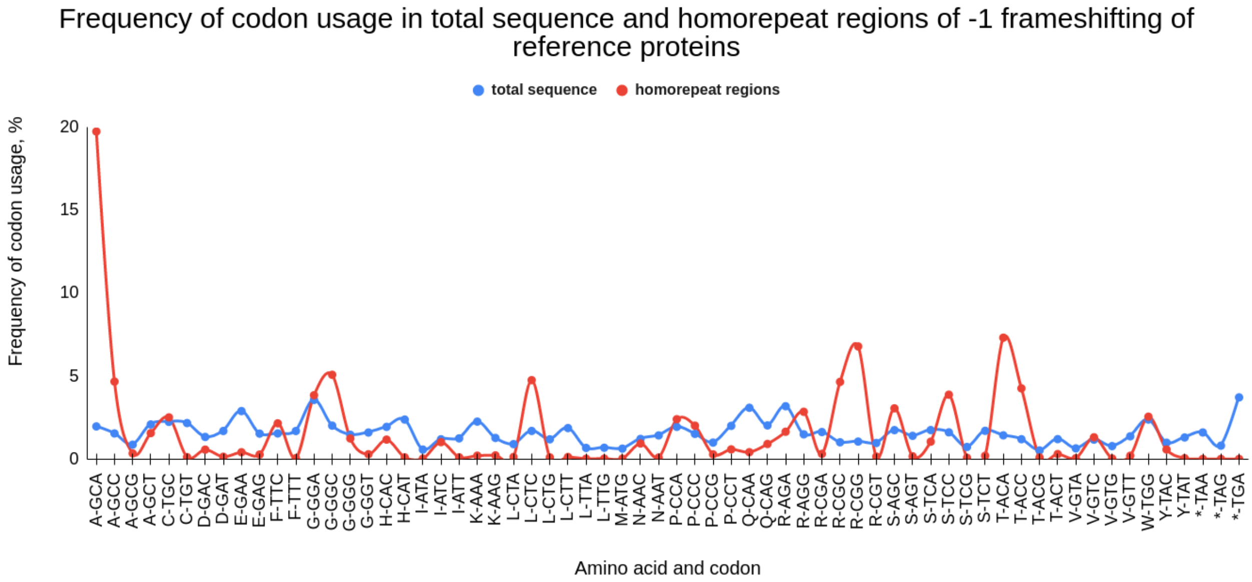


**Figure S6.** Codon usage in all -1 shifted reference proteins (blue) and -1 shifted homorepeats (red).

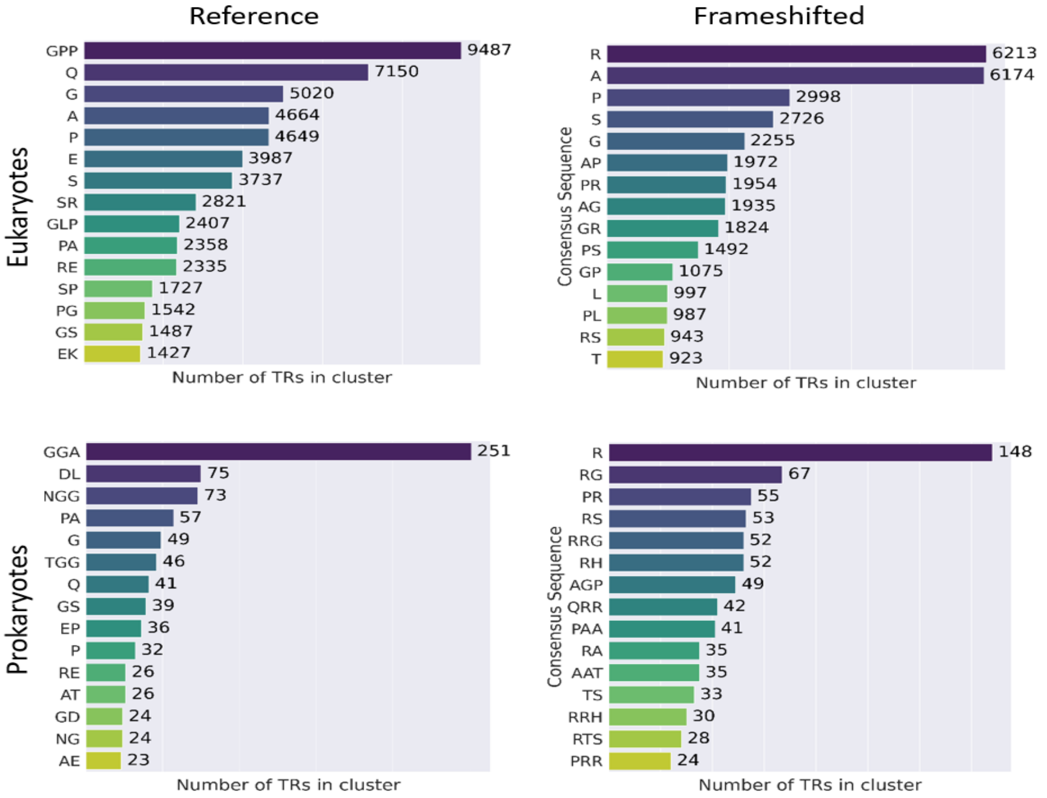


**Figure S7**. Top clusters of TRs in reference and frameshifted sequences of eukaryotic and prokaryotic organisms.

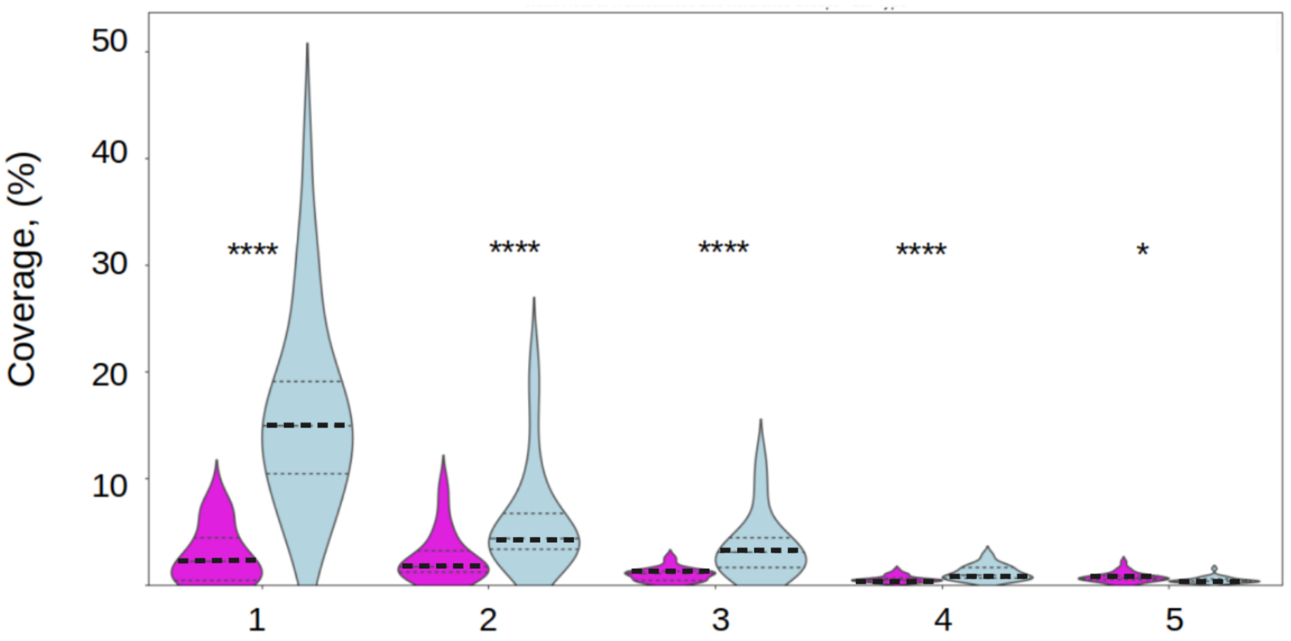


**Figure S8.** Coverage of exposed aggregation-prone regions (EARs) within TR regions of the proteins in eukaryotic organisms (frameshifted-magenta, reference-light cyan) predicted by ArchCandy. Statistical test performed by the two-sided T-test. **** and * mean p-value < 0.0001 and p-value < 0.05, respectively.

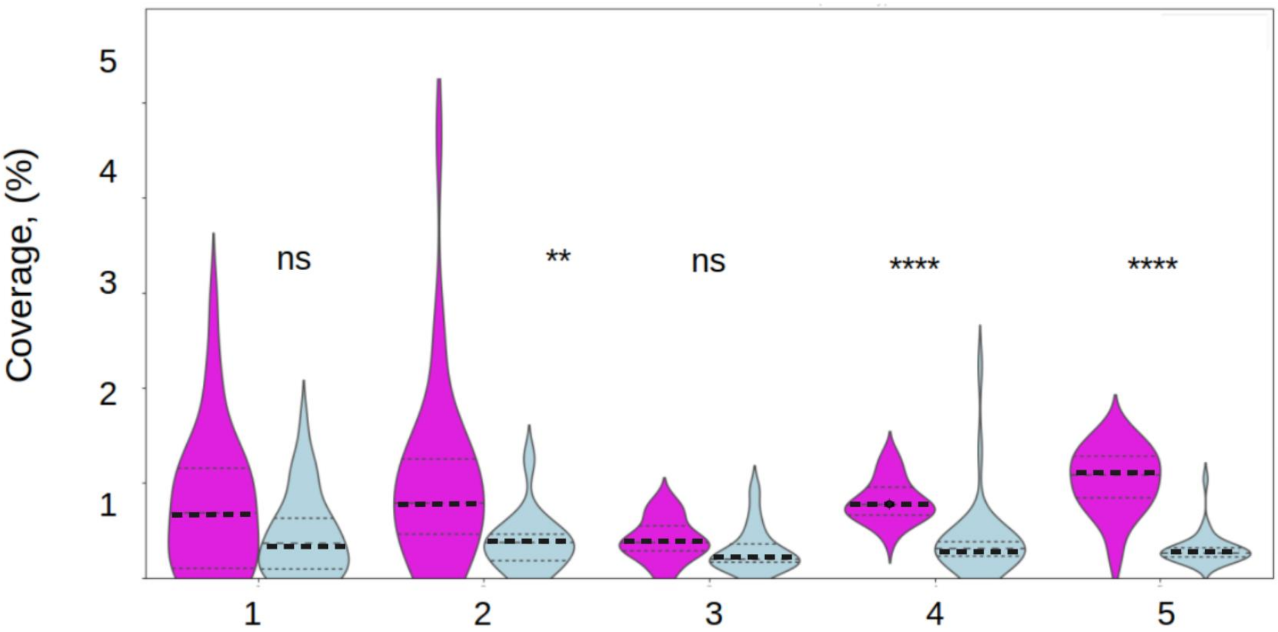


**Figure S9.** Coverage of EARs within TRs  of eukaryotic proteins (light cyan) and of their frameshifted sequences (magenta) predicted by TANGO. The observed variance performed by the two-sided T-test is statistically significant in TR groups, except Group 1 and Group 3. **** and ** mean p-value < 0.0001 and p-value < 0.01, respectively. ns means non-significant.
